## Supplemental Figures for "ST3Gal1 synthesis of Siglec ligands mediates anti-tumour immunity in prostate cancer"

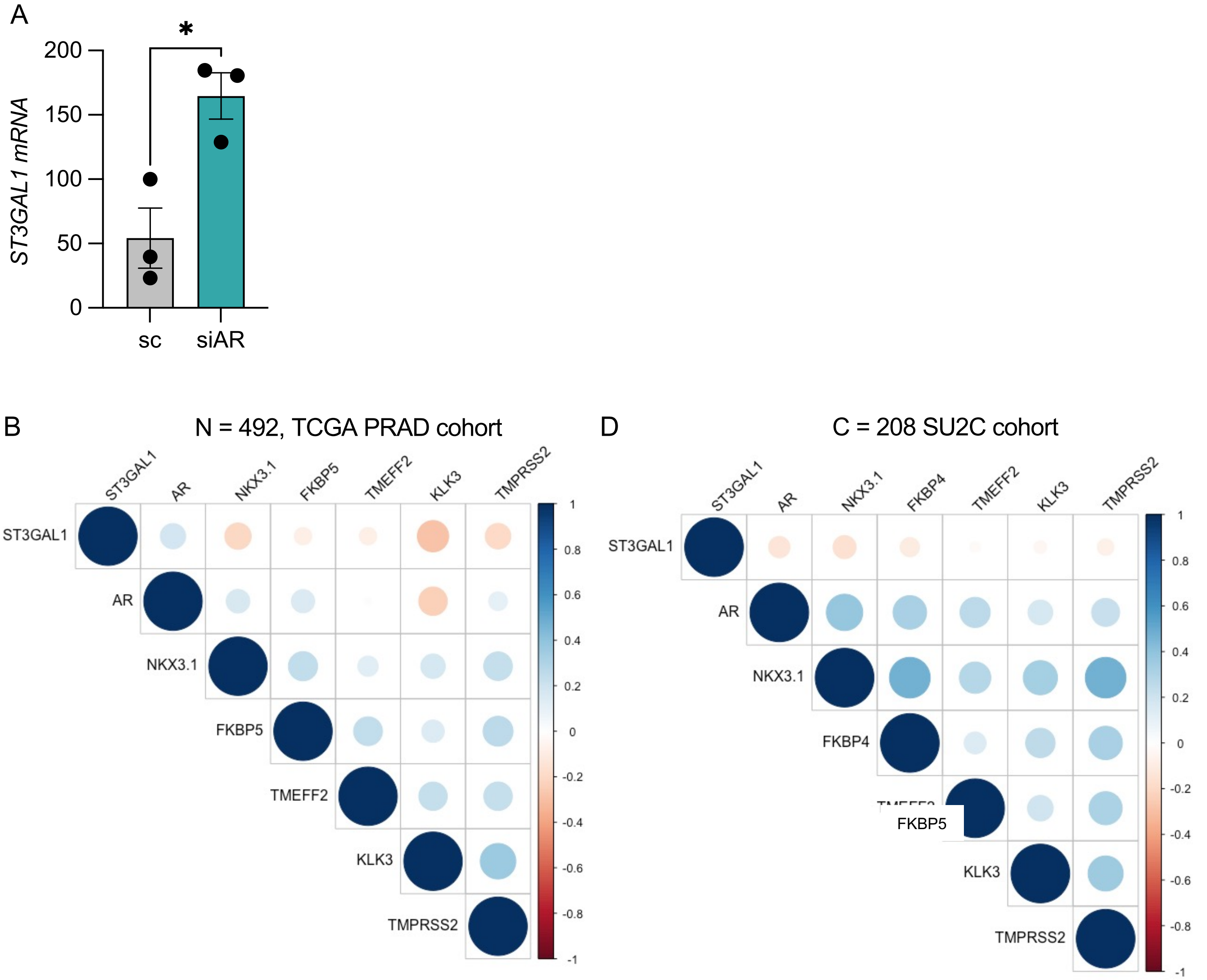


(A) Quantification of *ST3GAL1* mRNA by RT-qPCR in LNCaP cells following siRNA knockdown of full-length AR. (B) Correlation matrix correlogram showing *ST3GAL1* gene in prostate cancer patients (N=492). Pearson's correlation coefficient is shown from −1 (red) to 1 (blue). Only correlations with statistical significance of $p < 0.05$ are shown. The size of the circle is proportional to the correlation coefficients. (C) Correlation matrix correlogram showing *ST3GAL1* gene in CRPC patients (N=208). Pearson's correlation coefficient is shown from −1 (red) to 1 (blue). Only correlations with statistical significance of $p < 0.05$ are shown. The size of the circle is proportional to the correlation coefficients.

Supplemental Figure 1

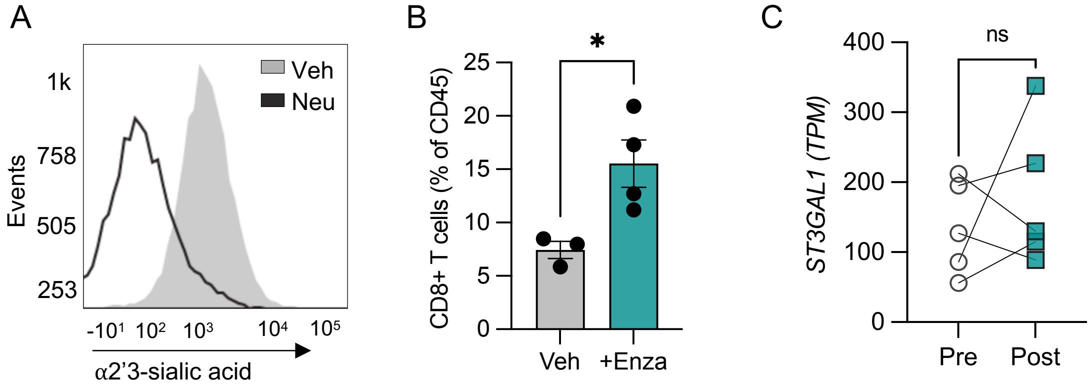


(A) Representative histogram of MAL-II lectin detection of α2'3-sialylation in LNCaP cells following following neuraminidase treatment. (B) Number of CD8+ T cells in TRAMP-C2 allografts following 20 mg/kg enzalutamide treatment for 7 days. Shown as a percentage of the total CD45+ population. (C) ST3Gal1 gene expression levels determined by RNA sequencing of match biopsies pre and post enzalutamide treatment (n=5).

Supplemental Figure 2

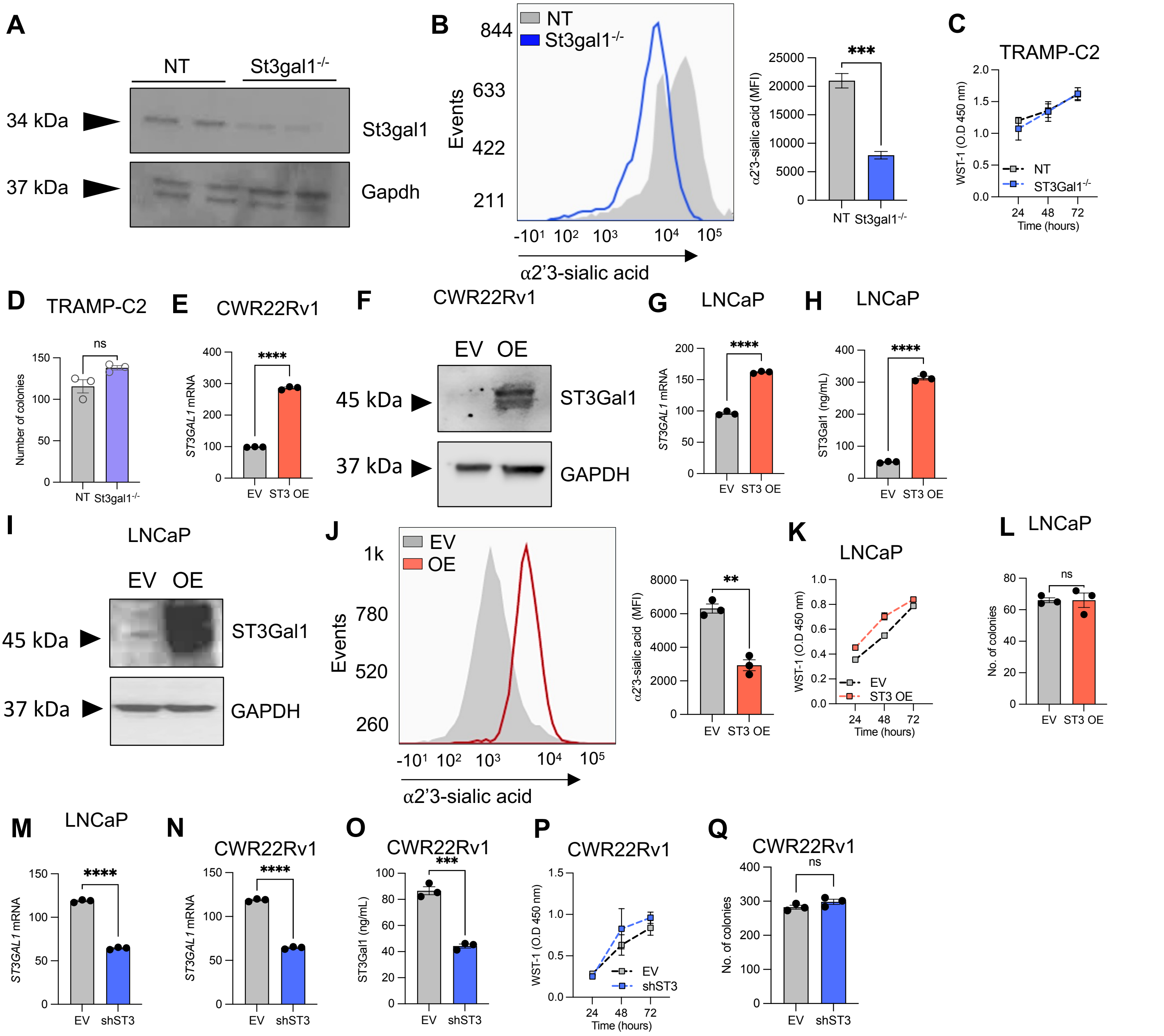


(A) Western blot for St3gal1 in non-targeting sgRNA TRAMP-C2 cells and *St3gal1*$^{-/-}$ cells. (B) MAL-II lectin detection of α2'3-sialylation in sgRNA TRAMP-C2 cells and *St3gal1*$^{-/-}$ cells by flow cytometry. (C) Cellular proliferation of NT and *St3gal1*$^{-/-}$ TRAMP-C2 cells quantified by WST-1 assay. Absorbance was read at 450 nm and normalised to background absorbance. (D) Colony forming ability of NT and *St3gal1*$^{-/-}$ TRAMP-C2 cells measured using a colony forming assay. Graph shows number of colonies formed. (E) RT-qPCR mRNA levels of *ST3GAL1* in empty vector (EV) and *ST3GAL1* overexpression (OE) lentiviral transduced CWR22Rv1 cells. (F) Western blot for ST3Gal1 in EV and OE CWR22Rv1 cells. (G) RT-qPCR mRNA levels of *ST3GAL1* in EV and *ST3GAL1* OE lentiviral transduced LNCaP cells. (H) Protein levels of ST3Gal1 in EV and *ST3GAL1* OE lentiviral transduced LNCaP cells determined by ELISA. (I) Western blot for ST3Gal1 in EV and OE LNCaP cells. (J) MAL-II lectin detection of α2'3-sialylation in EV and OE LNCaP cells by flow cytometry. (K) Cellular proliferation of EV and OE LNCaP cells quantified by WST-1 assay. Absorbance was read at 450 nm and normalised to background absorbance. (L) Colony forming ability of EV and OE LNCaP cells measured using a colony forming assay. Graph shows number of colonies formed. (M) RT-qPCR mRNA levels of *ST3GAL1* in EV and sh*ST3GAL1* lentiviral transduced LNCaP cells. (N) RT-qPCR mRNA levels of *ST3GAL1* in EV and sh*ST3GAL1* lentiviral transduced CWR22Rv1 cells. (O) Protein levels of ST3Gal1 in EV and sh*ST3GAL1* lentiviral transduced CWR22Rv1 cells determined by ELISA. (P) Cellular proliferation of EV and sh*ST3GAL1* CWR22Rv1 cells quantified by WST-1 assay. Absorbance was read at 450 nm and normalised to background absorbance. (Q) Colony forming ability of EV and sh*ST3GAL1* CWR22Rv1 cells measured using a colony forming assay. Graph shows number of colonies formed.

Supplemental Figure 3

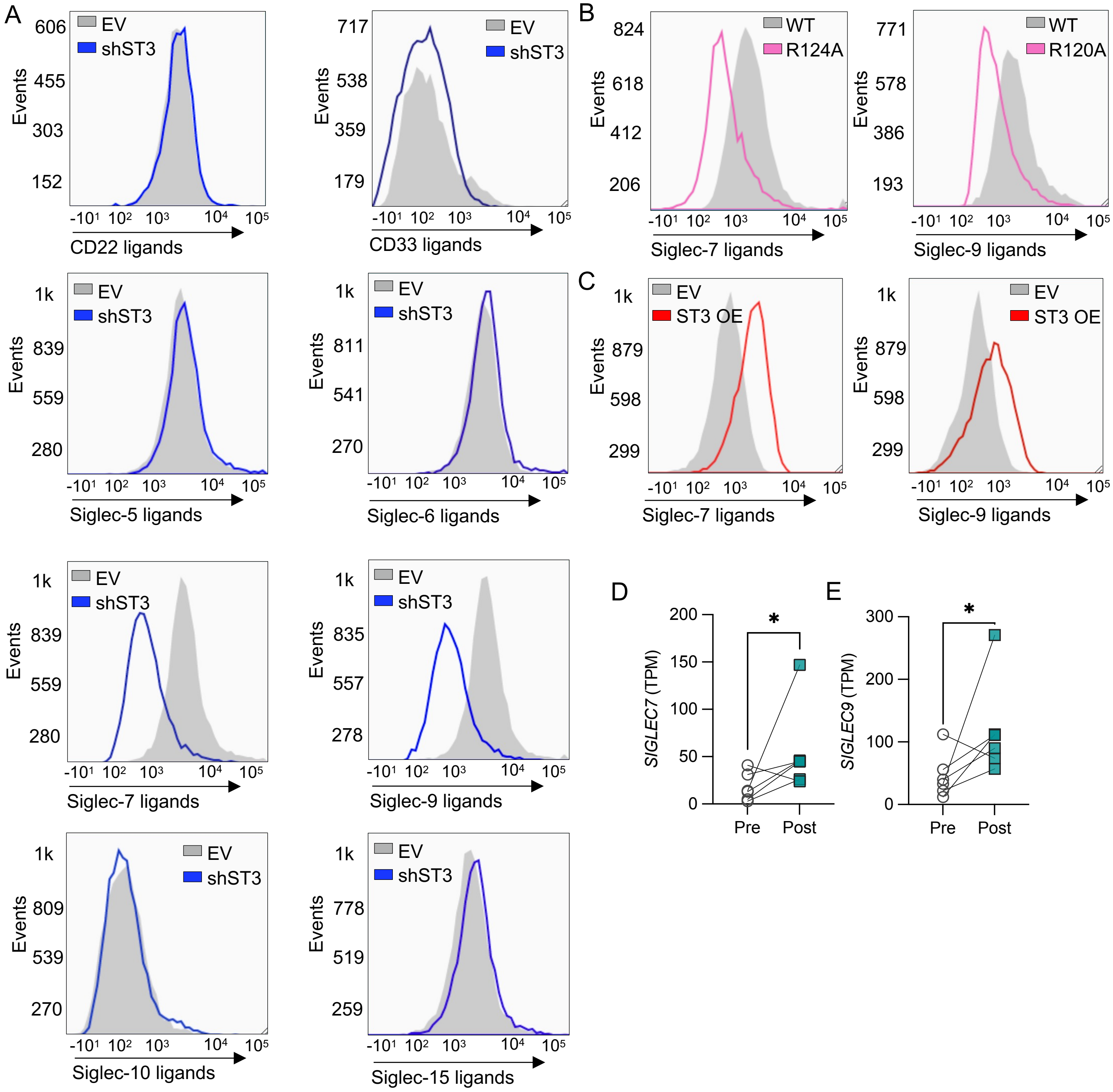


(A) Representative histograms for Siglec-Fc quantification of siglec ligands on empty vector and sh*ST3GAL1* LNCaP cells as determined by flow cytometry. (B) Representative histograms for engineered Siglec-9-Fc. WT Siglec-Fc is compared with an R120A/R124A mutant with reduced Siglec binding capacity. (C) Representative flow cytometry histograms for Siglec-7hFc and Siglec-9hFc quantification of siglec ligands on empty vector and ST3Gal1 overexpressing CWR22Rv1 cells (D-E) *SIGLEC7* and *SIGLEC9* gene expression levels determined by RNA sequencing of match biopsies pre and post enzalutamide treatment (n=6).

Supplemental Figure 4

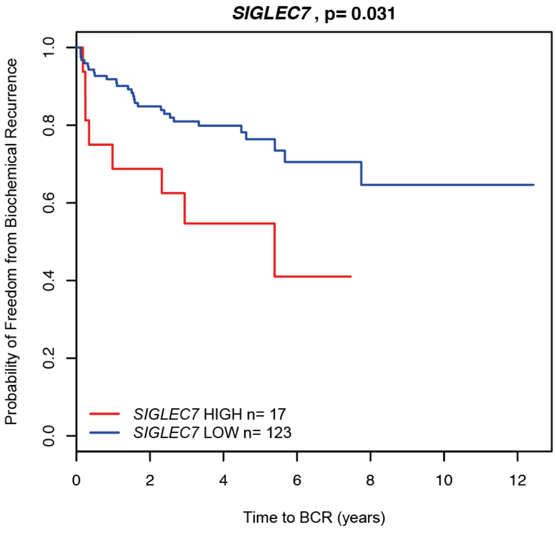


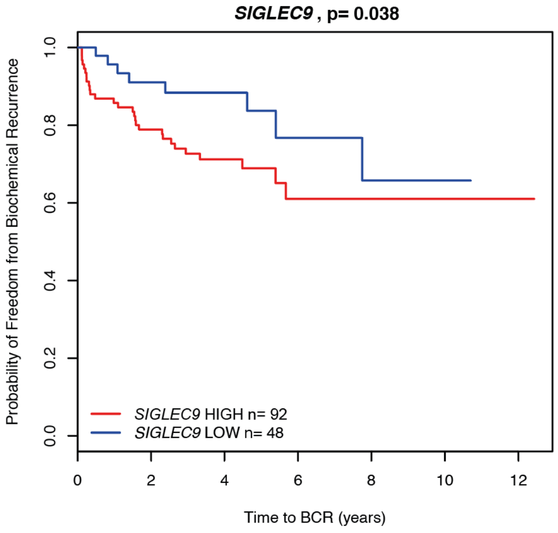


C

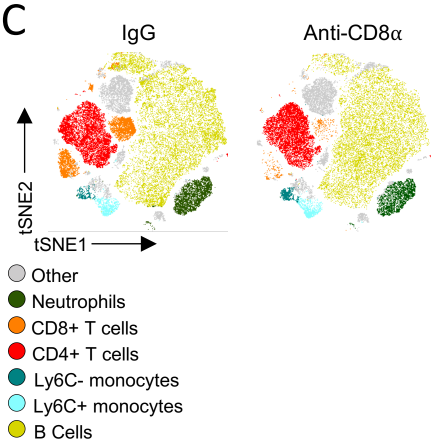


(A-B) Kaplan–Meier plot showing Time to biochemical reoccurrence for prostate cancer patients stratified based on low or high *SIGLEC7 and SIGLEC9 gene expression. Analysis performed on the MSKCC dataset accessed through camcAPP.* *(C)* t-distributed stochastic neighborhood embedding) tSNE maps of flow cytometry analysis of circulating immune populations in selective depletion TRAMP-C2 subcutaneous allografts treated with IgG or anti-CD8.

Supplemental Figure 5
